# Artificial selection of communities drives the emergence of structured interactions

**DOI:** 10.1101/2021.12.13.472438

**Authors:** Jules Fraboul, Giulio Biroli, Silvia De Monte

## Abstract

Species-rich communities, such as the microbiota or microbial ecosystems, provide key functions for human health and climatic resilience. Increasing effort is being dedicated to design experimental protocols for selecting community-level functions of interest. These experiments typically involve selection acting on populations of communities, each of which is composed of multiple species. Numerical simulations explored the evolutionary dynamics of this complex, multi-scale system. However, a comprehensive theoretical understanding of the process of artificial selection of communities is still lacking. Here, we propose a general model for the evolutionary dynamics of communities composed of a large number of interacting species, described by disordered generalized Lotka-Volterra equations. Our analytical and numerical results reveal that selection for total community abundance leads to increased levels of mutualism and interaction diversity. Correspondingly, the interaction matrix acquires a specific structure that is generic for selection of collective functions. Our approach moreover allows to disentangle the role of different control parameters in determining the efficiency of the selection process, and can thus be used as a guidance in optimizing artificial selection protocols.

---

Artificial selection has been used for millennia to steer plant and animal characters towards target phenotypes. Recently, it is attracting a lot of interest as a way to control and tune ecosystem services and functions, which are emergent properties of biological communities formed by many different species [1]. Particularly interesting in this respect are microbial communities that dispense highly relevant functions, contributing to human health [2] as well as to global biogeochemical cycles [3]. The widespread application of such an approach is nonetheless hampered by the large number of parameters that have potential bearings on the efficiency of the selection protocol, and that must be critically evaluated in designing these experiments [4].

Numerical simulations of simplified models have started exploring how selection for a collective function affects community composition [5–10]. Alternative experimental designs and system parameters have thus be shown to affect the efficiency of the selection process. Given the huge space of possible experimental choices and of interaction types, a fundamental problem is how to asses the robustness of simulation results and use them to optimize selection protocols.

Thorough studies of communities composed of two-species helped identifying key processes involved in artificial selection of communities, and pointed out how competition among composing species may be overcome in attaining collective functions [11, 12]. In particular, when community ecology was modelled by two-species competitive Lotka-Volterra equations, evolution of a specific community composition relied essentially on modifications of interspecific interactions [13]. In order to complement these studies, we consider complex communities composed of a large number of species, and develop analytical methods to address the statistical features of the evolutionary dynamics of such complex ecosystems. Approaches from statistical physics have been largely used to address the ecology of *species-rich* communities [14–19]. Generalized Lotka-Volterra equations with random interactions are an important null model for community ecology and a first step towards more realistic descriptions of the dynamics of complex ecosystems. Here, we propose and analyze a simplified model that tackles in general terms the effect of collective-level selection on the structure of interspecific interactions in species-rich communities. We study the evolution of collective functions in the limit when the ecological and evolutionary time scales are separated, where novelty is provided by mutations that affect collective functions. We derive a general equation describing the change along an evolutionary trajectory of the species interaction matrix. We show that selection for increased community size imprints a low-dimensional structure on such matrix, which results in the emergence of a global mutualistic term akin to collective cross-feeding. Our analytic results explain the effect of the number of replicate communities, community richness and diversity, nature of ecological interactions and magnitude of the mutational steps in determining the speed and attainability of a given target function upon which selection acts. This analysis reveals that community-level selection creates structure in interactions by progressively evolving a complex, structured matrix from an initially featureless one.

## MODEL FOR SELECTION OF SPECIES-RICH COMMUNITIES

We model a population of *n* communities that undergo cycles of ecological growth, selection and reproduction, as explained in Fig. 1. The ecological dynamics within a cycle is described as a function of a continuous time variable *t*. Reproduction occurs via monoparental seeding of the next community generation (’propagule’ reproduction [5, 8]). The successive cycles, or community generations, are indexed with a discrete variable *τ*. Selection is applied by letting the probability that a community reproduce depend upon a collective function, evaluated at *t* = *T*, the duration of one generation. The evolutionary dynamics that we aim to describe consists in the change of the community composition across multiple generations. For simplicity, we assume that mutations only occur in newborn communities, so that within one collective generation species abundances are only ruled by the ecological dynamics.

**Figure 1.**
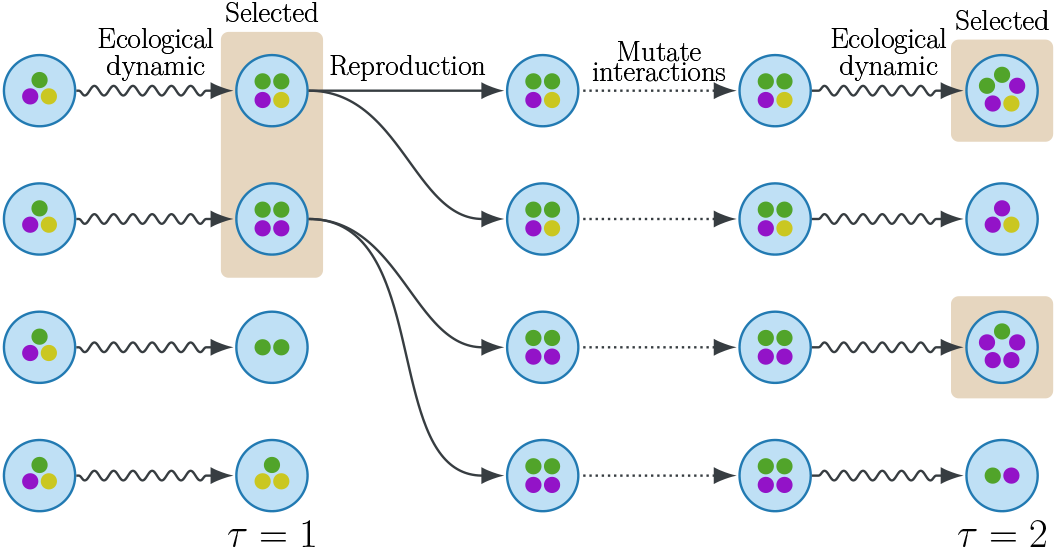
Structure of the model for the evolution of species-rich communities. Each community in a population of *n* (here, *n* = 4) communities is represented by a circle and is composed of a set of individuals (represented by the dots), belonging to different species (represented by colour), initially sampled by a same metacommunity. The *m* = 2 communities with largest total abundance (the number of individuals) are selected for reproduction. Newborn communities are generated by copying the state (vector of species abundances), but modifying the parameters of the interactions among species, as detailed in the text. In the course of a community generation, these changes can result in ecological variation of community composition and of its selected function.

In the following, we first detail the model for the dynamics of a single community within one generation, and then the rules for community reproduction and mutation.

*Ecological dynamic*. Each of the *n* communities is composed of *S* species with continuous abundances (*N*_*i*_)_*i*=1,…,*S*_, whose variation is described by the generalized Lotka-Volterra equations [20]:

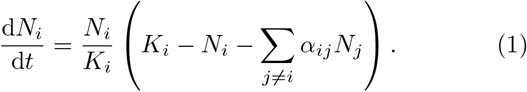

The constants *K*_*i*_ are the carrying capacities and the interaction coefficients *α*_*ij*_ represent the effect of species *j* on the growth of species *i*.

### Initial species interactions

Following May’s disorder approach [21], we choose initial communities with random interactions. Specifically, the coefficients *α*_*ij*_ are drawn, as in Bunin, from a normal distribution of parameters:

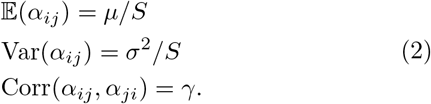

Here, *μ* represents the total interaction strength faced by one species from all of its partners, whereas *σ* measures the diversity of interactions. The parameter *γ* ∈ [−1, 1] determines the symmetry of the ecological interactions: competition and mutualism correspond to *γ* = 1 whereas exploitative interactions like predator-prey and parasitic interactions are characterized by *γ* = −1.

We initialize communities by sampling interactions in the region of parameters (*μ, σ*) where the system has a unique, stable coexistence equilibrium (see Supplementary Section 1), that is independent of the initial community composition [15]. The transient dynamic leading to such an attractor depends however on the initial state of the community, and can have important evolutionary implications [8]. For this reason, the duration *T* of one generation is assumed to be large, so that the community approaches equilibrium.

### Community-level selection and reproduction

Communities are ranked according to a single collective function. We will mostly focus on the total community abundance *N*_*T*_ = Σ_*i*_ *N*_*i*_(*T*). The *m* communities that rank best at the end of one generation are chosen for reproduction, and the rest discarded (Fig.1). When an offspring community is born, it acquires the same composition of the parent community. In the absence of variation in the community parameters, this guarantees that community functions are perfectly inherited.

### Community-level mutations

For evolution by natural selection to occur at the level of communities, there must be variation in the collective function [22]. In our model, variation is replenished at each community generation by changes in the interaction matrix, called ‘community-level mutations’. In order for mutations not to bias a priori the change of the trait under selection, mutations are defined so that they do not alter, in expectation, the mean and variance of the interaction matrix. Single realizations nonetheless differ in the collective function, producing the variation selection acts upon. To this purpose, we write the interaction matrix at generation *τ* as:

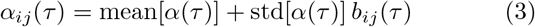

where:

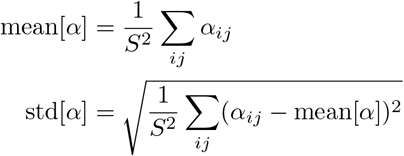

are the empirical mean and standard deviation of the matrix *α*, and the reduced matrix *b* has empirical mean 0 and empirical variance 1.

We define the mutated interaction matrix as:

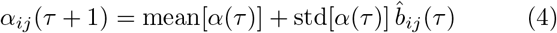

with:

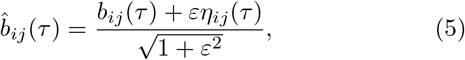

where *η*(*τ*) is a Gaussian random matrix of expected value 0, variance 1 and symmetric correlation *γ*. Therefore, the interaction matrices at two successive generations have the same probability distribution. In the absence of selection, thus, community function will evolve by *neutral drift*.

## RESULTS

We now present and discuss the salient features of the evolutionary dynamics of the model previously introduced. We first show numerical results and then addressed them theoretically. The numerics is performed for initially mostly competitive interactions and randomly distributed carrying capacities, as detailed in the Materials and Methods.

As observed in past numerical studies [5, 6, 23, 24], we find that in *response to selection*, communities evolve so as to improve the desired collective function (Fig. 2). Such increase gradually accelerates, and the ecological dynamics is eventually pushed in a region where some abundances diverge (This divergence is a well-known pathology of the Lotka-Volterra equations that can be corrected by choosing a saturation stronger than quadratic [25]).

**Figure 2.**
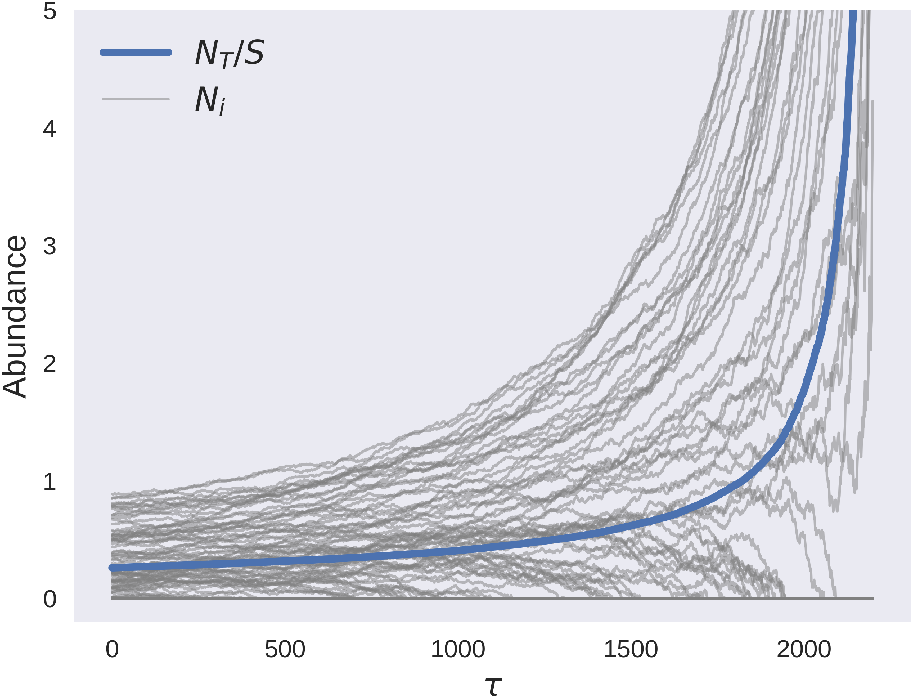
Changes of species abundance along an evolutionary trajectory. Selection for increased total abundance leads to an increase in the abundances of most species (grey lines), and, as a consequence, of the average abundance *N*^*T*^ */S* (blue line). See Material and Methods for the details of the numerical simulations.

The observed improvement of community function derives from *changes of the interaction matrix α*, that also reflect on its empirical statistics *μ*(*τ*) = mean[*α*(*τ*)]/*S* and 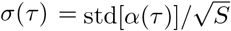 As shown in Fig. 3 A, the mean decreases, indicating that interactions become – on average – progressively more mutualistic. At the same time, their variance increases, so that interactions within the community become more diverse.

**Figure 3.**
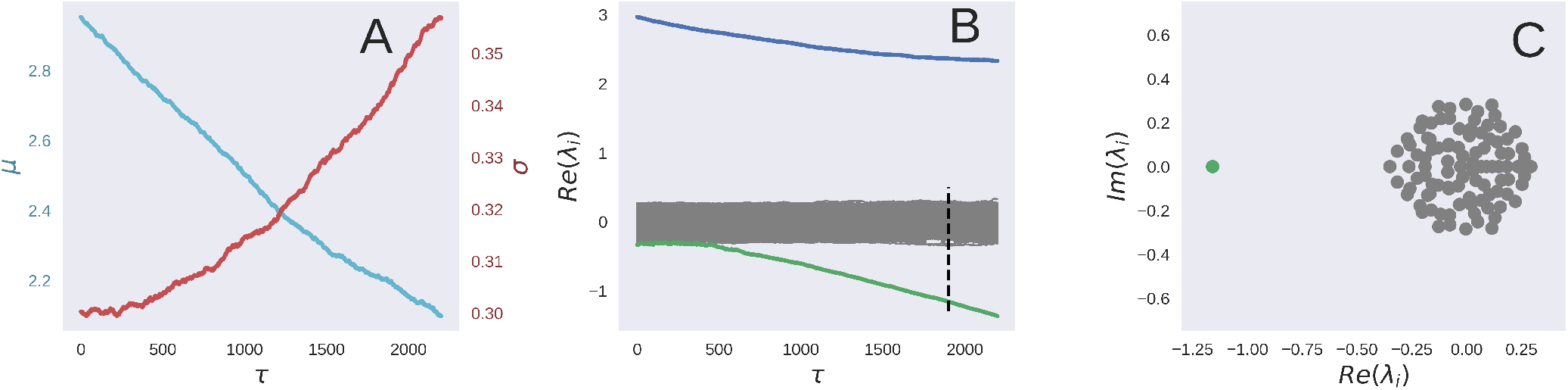
Changes of the statistics of the interaction matrix along an evolutionary trajectory. The interaction matrix *α* of the best community evolves so that the average interaction strength decreases linearly in time (A, cyan), while its variance increases (A, red). Such changes correspond to a modification in structure, manifest in the spectrum of its eigenvalues. The change of their real part across community generations (B) reveals the appearance of an isolated negative real eigenvalue (green) as well as the decrease of the eigenvalue associated to *μ* (blue). A zoom of the spectrum in the complex plane (C) at generation *τ* = 1900 represented by the dotted line in (B) reveals that, apart from the emergence of this mutualistic collective mode, the matrix retains its initial random structure characterized by a circular law for the eigenvalues.

Analytical results obtained for disordered communities can help to rationalize these findings. Indeed, for random matrices defined by equation 2, the total population size *N*_*T*_ is purely a function of *μ* and *σ*. Thus, one could envision selection as a process in which the empirical moments of *α* change across community generations so as to climb the gradient of the fitness function *N*_*T*_ (*μ, σ*) (reproduced from [15] in Supplementary Fig. 1). This, however, is not what happens: the evolutionary trajectory of the community function *N*_*T*_ (*μ*(*τ*), *σ*(*τ*)) deviates from the gradient-climbing process predicted for a random matrix with the same moments (Fig. 4). Hence, evolutionary dynamics cannot be described as a change of global features characterizing the interactions. One needs to dwell on the evolution of the fine-scale properties of the interaction matrix.

**Figure 4.**
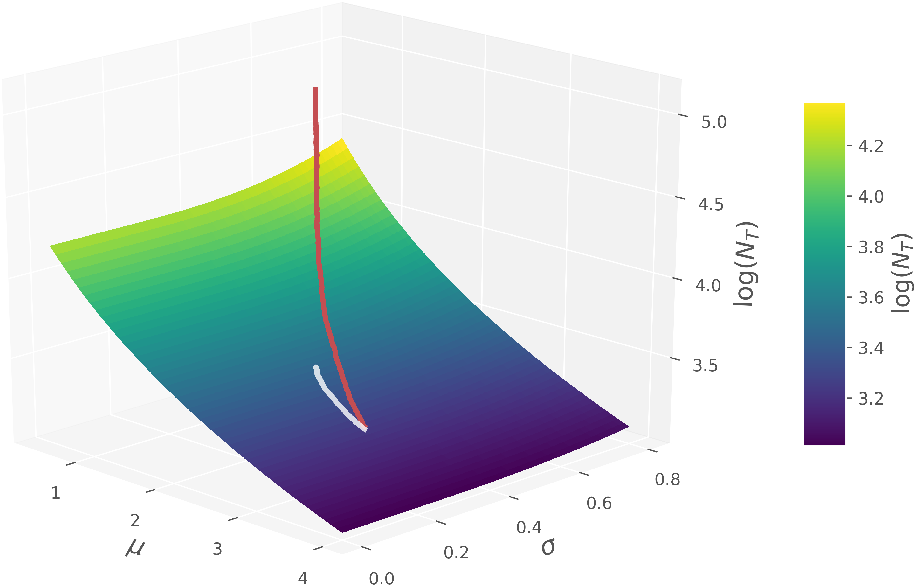
Purely random interactions cannot explain the evolution of total community abundance. Variation of the interaction moments *μ*(*τ*), *σ*(*τ*), and of the total abundance log(*N*^*T*^ (*τ*)) (red line) along an evolutionary trajectory. The abundance of a random interaction matrix (equation 2) with moments *μ, σ* (surface) is plotted for comparison. The white line is the predicted total abundance if the matrix of moments *μ*(*τ*), *σ*(*τ*) was completely random, indicating that along the trajectory the matrix becomes progressively structured.

As we show below, *selection imprints a structure* on the interaction matrix *α*, that can be characterized by its eigenvalues. The spectrum of the initial random interaction matrix is, in the complex plane, a circle of radius *σ* centered in the origin [26], plus an isolated positive eigenvalue (blue in Fig. 3 B) of magnitude *μ*. The initial effect of selection is to reduce this value. After some time, however, an isolated *negative* eigenvalue *λ* (green in Fig. 3 B and C) emerges from the circle and detaches from it linearly in time. Apart from this isolated component, along an evolutionary trajectory the matrix retains its randomness, and the circle of eigenvalues changes only slightly its radius. Selection adds to the random part a new rank-one term which can be written in terms of the eigenvectors relative to the isolated eigenvalue *λ*.

The imprinted structure that emerged along the evolutionary trajectory can be visualized by displaying the corresponding entries of *α* for early and late stages of community evolution (Fig. 5, where species are ordered in terms of their carrying capacity from larger to smaller). At the beginning, there is no visible structure, except the diagonal that has zero entries by construction. After 2000 generations, species who have become more mutualistic are mostly those that initially had higher carrying capacity. This is a direct manifestation of the emergence of the isolated eigenvalue, as we find that its eigenvectors are correlated to both **K** and to the equilibrium abundances **N** (see Supplementary Fig. 3).

**Figure 5.**
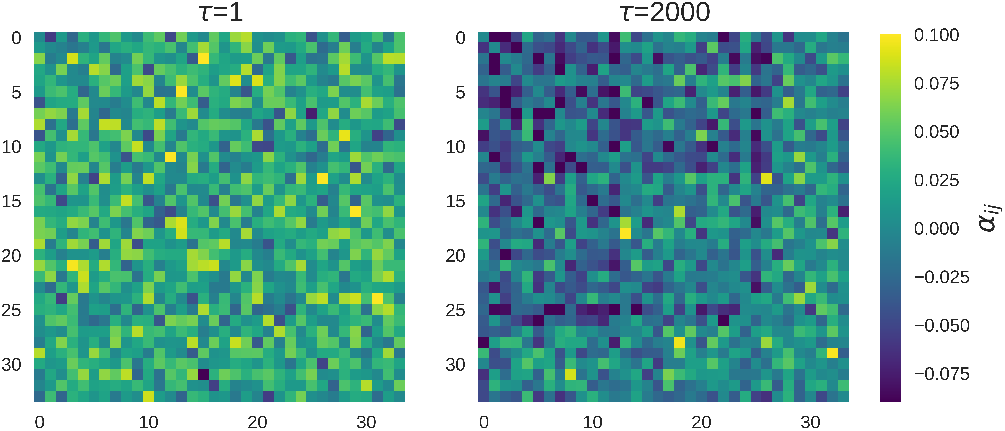
Evolution of the interaction matrix. Coefficients of the interaction matrix *α* with rows and columns sorted by decreasing carrying capacities at generations 1 (left) and 2000 (right) for the same simulation as Fig. 2. Only the species that have positive abundance at generation 2000 are shown.

Simulations realized for a number of different parameter values and for other target functions (Supplementary Section 10) suggest that the phenomena illustrated above are general. Next, we introduce a *theoretical framework* to explain how the phenomena illustrated above by numerical simulation emerge and why they should be expected to hold generically. Analytical results, illustrated here for asymmetric interactions (*γ* = 0) and detailed in Material and Methods and Supplementary Information, moreover allow us to disentangle the respective role of all the system’s parameters in determining the efficiency of the artificial selection process.

Given a community with interaction matrix *α*(*τ*) and equilibrium **N**(*τ*) at a given generation *τ*, we aim to characterize the interaction matrix *α*(*τ* + 1) of the selected offspring community – that providing the largest total abundance at equilibrium. Mutations of *α*(*τ*) are, for *ϵ* ≪ 1, equivalent to small random perturbations of the carrying capacities, so linear response theory provides the corresponding change induced on the equilibrium abundances. The total abundance of each of the *n* communities is therefore modified by a random contribution that we can fully characterize. By singling out the largest contribution, i.e. the largest among several independent random variables [27], we show in the SM that selection induces the following change in total abundance *N*_*T*_ (*τ*):

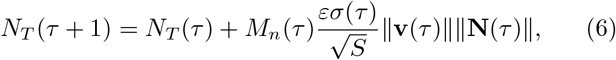

where **v**(*τ*) is a nonlinear function of the interaction matrix *α* (*τ*): v^*^ (*τ*) = (𝕀 * + α * (*τ*)^T^)^*−*1^**1** (the asterisk indicates that only extant species are considered) and *ν*_*i*_ = 0 for extinct species. This vector also measures how the total abundance at equilibrium varies upon changing the carrying capacities: 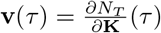. The random variable *M*_*n*_(*τ*) (drawn independently at each generation) follows the statistic of the maximum value of *n* Gaussian variables, with expected value 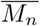 (see the distribution of *M*_*n*_ in Supplementary Fig. 2).

Equation 6 implies that the total community abundance increases on average along an evolutionary trajectory, as the product of the norms is always positive and *M*_*n*_ has a positive expected value for *n >* 1. However, when the number of communities is too small, it can also transiently decrease, thus breaking the alignment between selection and community response.

Changes in total abundance are ultimately based on the evolution of the interaction matrix. As detailed in the SI, its change across one collective generation can be decomposed in a directional term – contributing to the evolution of *N*_*T*_ – and its complement *B*_*ij*_, that acts as a random fluctuation. The interaction between any two species *i* and *j* thus evolves according to:

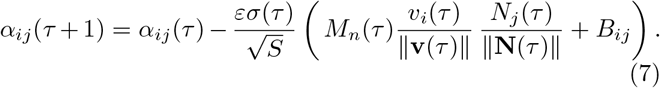

This expression has a simple interpretation: among the random mutations of the interaction matrix, only matter those in the special direction associated to the target function *N*_*T*_. The selected community is hence the one having the largest random Gaussian contribution associated to such direction. Equation 7 (or its generalization Supplementary equation 22 for *γ* ≠ 0) means that species are not all equivalent in the face of selection: species whose potential variation contributes more to the function (those with larger *v*_*i*_) and with larger equilibrium abundance will face a larger decrease in the interaction strength, as pointed out in Fig. 5.

Equations 6 and 7 apply to any initial interaction matrix (not only random ones) and allow us to draw general conclusions on how *speed and direction of evolutionary change* depend on the numerous parameters of the system.

As could be intuited, evolution is faster when selection screens a larger number of communities, since the expected value 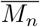 is an increasing function of *n*. When only one community is considered, on the other hand, the total abundance and the interaction matrix undergo unbiased stochastic changes (see Supplementary section 7), as *M*_1_ is Gaussian with zero mean. Under these conditions, collective functions cannot be selected and evolve by community-level drift. However, increasing the number of communities may not always be the key to success. The growth of 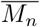 with *n*, indeed, scales as 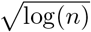, which increases slowly for large *n*, so that transition to high community throughput may be of little avail to speed up evolution.

Other parameters can be changed so as to improve the efficacy of community selection. The variation of the interaction matrix, thus of the selected function, across one community generation occurs on a time scale 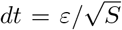. Thus, in our model evolution has faster pace in communities with a smaller number of species and for larger mutational steps.

Equation 7 (equation 22 for general *γ*) are non-linear recursive matrix equations that can’t be exactly solved in general. A complete solution can however be obtained in the limiting case of small variability of interactions *σ* ≪ 1 (see Supplementary section 6). The effect of selection is here simply to add a global negative term to the dynamics of the average interaction strength and results in interactions becoming progressively more mutualistic (Supplementary equation 30). A phenomenon discovered in random matrix theory and called BBP phase transition sheds light on the origin of the structure observed in simulations. Such transition is characterized by the emergence of an isolated eigenvalue *λ* when a strong enough rank one term is added to a random matrix. Fully characterized in random matrices [28], the BBP transition found applications in computer science, data sciences, neurosciences and physics [29–33]. An identical phenomenon takes place in our case, as shown in Fig. 3 C, and leads to an isolated eigenvalue (with left and right eigenvectors **q** and **r**). Analogous to the BBP phenomenon, the emergence of the isolated eigenvalue is the consequence of the addition of rank-one directional contributions in eq. 7 (whereas the random Gaussian contributions *B*_*ij*_ do not modify the initial random structure). Unlike the BBP transition, however, these contributions change in time, so that the exact moment when the transition occurs is only predictable when the heterogeneity of interactions is small.

A full theory can be nonetheless obtained when some community features can be observed along an evolutionary trajectory. By leveraging the maximum entropy method method developed in Barbier *et al*. we can characterize the interaction matrix from the mean interaction *μ*, the equilibrium abundances **N** and the carrying capacity vector **K**. For large *S*, this matrix can be written (see Supplementary section 9):

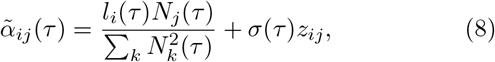

where *l*_*i*_(*τ*) = *K*_*i*_ − *N*_*i*_(*τ*) − *μ*(*τ*)*N*_*T*_ (*τ*)*/S* and *z*_*ij*_ is a Gaussian random matrix of zero mean and unit variance.

Remarkably, this expression provides a very accurate description of the evolutionary dynamics of the interaction matrix (see Supplementary Fig. 8). In particular, the isolated eigenvalue and the associated eigenvectors show a perfect match. Within this framework, the interaction matrix at evolutionary time *τ* is indeed the initial one (restricted to extant species) plus a time-dependent rank-one contribution. BBP theory therefore implies that the effect of evolutionary dynamics on the spectrum is to: (i) shrink the circle of eigenvalue (due to *σ*(*τ*)*z*_*ij*_ and whose radius is equal to *σ*(*τ*)), (ii) induce a BBP transition with a lower eigenvalue popping out when ‖**l**(*τ*)‖ > *σ*(*τ*) ‖**N**(*τ*)‖. These theoretical results provide a quantitative explanation of our numerical findings and suggests that it should be possible to identify the emergence of the isolated eigenvalue without measuring every pairwise interaction coefficient – something that would be unreasonably demanding in actual experiments.

The equations for the evolution of the interaction matrix can be generalized to other selection targets such as linear combinations of abundances at equilibrium (see Supplementary section 10). The emergence of a rank-one structure is hence not specific to the increase of the total abundance. This is supported by the fact that the our arguments are valid for any selection targets depending only on the abundances at equilibrium.

## DISCUSSION

This study is devoted to identifying key and general features of the evolutionary dynamics in species-rich communities under a scheme that is commonly used for artificial selection of collective functions. We showed that the interaction matrix evolves in response to selection for total abundance, and that it results generically in interspecific interactions becoming progressively less competitive. We interpret this as the evolution of facilitation, similar to what was observed in a two-species model [13]. At the same time as the average strength of interspecific interactions decreases, they become more variable. Notably, the evolutionary process imprints a structure on the interaction matrix. The key to this structure is an isolated eigenvalue, which emerges as a ‘collective mode’ that positively impacts the abundances of all species. In the analytical description, this corresponds to an order-one perturbation of the interaction matrix, that otherwise retains its original, disordered nature. If we consider the Lotka-Volterra model as a limit case of the MacArthur equations, which describe not only species abundance, but also the resources they consume, the emergent facilitation term can be viewed as an effective global cross-feeding. This result is not specific to selection acting on total abundance, but seems to hold for any function of the abundances at equilibrium. Our finding resemble phenomena observed in other domains where interactions between degrees of freedom are adjusted dynamically to lead to a specific collective property, such as a lower ground state energy in spin-glasses and learning in neural networks [35–37]. The ubiquity of low-rank perturbations raises the question of if and when selection can produce the emergence of more complex structures.

We chose to analyze an idealized model in order to achieve analytical tractability. Although it can be argued that disordered models are an oversimplification of real communities, they provide null expectations for collective properties. Moreover, the actual strength of ecological interactions is unknown in most microbial communities. Given this state of affairs, and that not all detailed properties of the interactions are expected to matter in shaping the general behaviors of ecosystem, the statistical approach appears a valuable method for identifying general prescriptions relevant even in experiments [38, 39].

This model may be extended in several meaningful ways. Instead of modelling species interactions through direct effects, one could include explicitly the resources that are consumed or exchanged [5, 8, 40]. Given the equivalence of the Lotka-Volterra and MacArthur models when resource dynamics is much faster than the ecological one, we do not expect this extension to qualitatively affect the main results of our work. However, a formulation in terms of resource consumption would connect theoretical results to experiments exploring the metabolic foundations of ecological interactions in microbial communities [41, 42]. Especially, this may guide the choice of more realistic interaction matrices, such as sparse networks [43, 44], or networks with empirical biases [45].

Even maintaining random direct interactions, the model we considered could be explored in regimes where the perturbative approach is expected to break down. This would occur for instance when the ecological dynamics of the community does not attain an equilibrium because of transients [8], stochastic demographic fluctuations [17] or chaotic population dynamics [15, 16, 46]. All these processes may reduce the heritability of the community function and thus alter the evolutionary trajectory.

Finally, consistent with the idea that communities are Darwinian individuals [47], we chose mutations that would provide unbiased community-level variation in the target function. Such assumption allowed us to develop a null model that is independent of the details of the underlying community interactions. Collective-level mutations can be thought of as the result of multiple changes in species interactions that occurred during the lifetime of a community. More detailed descriptions of how sequential species-level mutations give rise to variation of the interaction matrix at the time of community reproduction – that is when the function is evaluated – may prove necessary for specific applications, and provide additional constraints, as observed for simpler models [13]. Furthermore, the model could be extended so as to include mutations of intra-species interactions via changes of the carrying capacities or speciation events that would increase diversity.

Communities are increasingly conceived as coherent units that provide collective-level functions, to the point to be attributed the status of ‘organisms’ [48, 49]. If this view can reflect the way ecological interactions produce a given population structure [50], it can go as far as identifying communities as full-fledged evolutionary units. In the latter case, how they are ‘scaffolded’ by physical compartmentalization and the establishment of community-level lineages, is all-important in determining the action of natural selection at the level of communities [51, 52]. We have modelled here the protocol commonly used in experiments of artificial selection [4, 7]. Considering that the collective level is the true center of interest for this process, moreover, we described mutations only for their effect on the community-level function under selection. Nested levels of reproducing units are widespread in the hierarchy of living beings. Our results might thus be relevant whenever selection on high-level functions bestows a structure on the interaction among heterogeneous constituent units, and contribute to understanding how integration across levels of organization is achieved.

## MATERIALS AND METHODS

### Description of the code

Numerical simulations were performed in python using the code accessible at https://github.com/jules-fbl/LV_community_selection. All the figures of the paper were obtained with a number of species *S* = 100, *m* = 1 selected community out of *n* = 10, a mutation strength *ε* = 0.02, an initial interaction matrix drawn from a Gaussian distribution of parameters *μ* = 3, *σ* = 0.3 and *γ* = 0 and random carrying capacities drawn uniformly between 0.5 and 1.5. The collective generation time was chosen to be *T* = 500 (with the exception of the first generation where a time *T* = 5000 was used in order to avoid the propagation of transient effects). This time is long enough for the mutated communities to approach their equilibrium abundances. To integrate the Lotka-Volterra equations, we used an integration scheme described in Supplementary section 11. We also imposed an abundance cut-off *N*_min_ = 10^*−*20^ below which species are deemed extinct.

### Derivation of recursive equation for the interaction matrix

We here explain the derivation of equation 7 in the case *γ* = 0. The complete derivation can be found in Supplementary section 10.

Let *α*_*ij*_ be the interaction matrix of a community at generation *τ*, of empirical mean and variance *μ/S* and 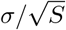 and let **N** be the associated abundances at equilibrium. After a mutational step, the interaction matrix becomes, at first order in *ε* :

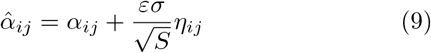

with *η* a Gaussian matrix of expected value zero and variance1 The mutation of *α* is equivalent to altering the carrying capacities by 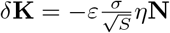.

We define the perturbation matrix as 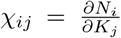. This matrix measures the effect of a small change in the carrying capacities on the abundances at equilibrium and is related to the interaction matrix through the identity *χ* = (𝕀^⋆^ + *α*^⋆^)^*−*1^ (see Supplementary section 2). Then if 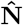 are the equilibrium abundances associated to the mutated matrix, the variation 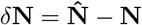 can be seen as a first order perturbation:

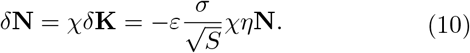

The induced variation *δf* of the total abundance is:

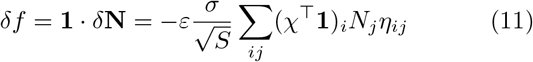

For *n* realizations of matrices (*η*_1_, …, *η*_*n*_) (one for each new-born community), we can find the distribution of the matrix that maximizes *δf* by using extreme value statistics (see Supplementary section 3):

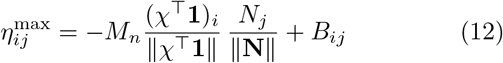

with *M*_*n*_ a random variable following the statistic of the maximum value of *n* Gaussian variables and *B* a Gaussian matrix. Then, the selected interaction matrix at generation *τ* + 1 is obtained by putting the expression of *η*^max^ in equation 9. Defining **v** = 𝕀^T^**1**, we get equation 7.

## Supporting information

Supplementary Informations

## ACKNOWLEDGMENTS

We thank Matthieu Barbier for insightful discussions. This research was partially supported by a grant from the Simons Foundation (N. 454935 Giulio Biroli). SDM was supported by the French Government under the program Investissements d’Avenir (ANR-10-LABX-54 MEMOLIFE and ANR-11-IDEX- 0001-02PSL).

