## Supplementary Informations for "Artificial selection of communities drives the emergence of structured interactions"

#### Contents


**1 Phase diagram** 2

**2 First order perturbation theory of Lotka-Volterra equations** 2

**3 Maximum over Gaussian samples** 3

**4 Mutation-Selection Process for any $\gamma$** 4

**5 Correlation of the eigenvector with the abundances at equilibrium** 5

**6 Limit of small $\sigma$ and large $S$** 5

**7 Community changes in the absence of selection** 7

**8 Evolution of species diversity** 8

**9 Comparison with synthetic interaction matrix** 9

**10 Generalizations** 9

**11 Numerical integration method** 9

### 1 Phase diagram

The phase diagram of the random Lotka-Volterra model (equations 12 of the main text) in the space of the mean $\mu$ and standard deviation $\sigma$ of the interaction matrix, for the case $\gamma = 0$, was derived in [Bunin, 2017] and is reproduced in Figure 1. In the unbounded growth phase, some population sizes diverge in finite time: this is a pathological feature of the Lotka-Volterra equations, that can only be corrected by complexifying the equations. The chaotic phase is characterized by a chaotic dynamics with multiple unstable equilibria. In the unique-equilibrium phase, the community converges, independently of the initial conditions, toward a unique ecological equilibrium where numerous species coexist. In light of this result, we decided to draw our initial interaction matrix in the unique-equilibrium phase as the ecological equilibrium is well defined. In the last two phases, a long-term value of the mean abundance can be computed, depending only on $\mu$, $\sigma$ and $\gamma$: the contour plot of the log of this mean value is represented in Figure 1. Being the total abundance proportional to the

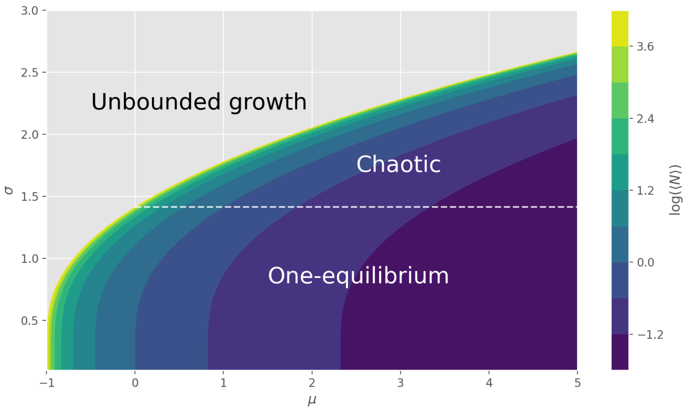


Figure 1: Phase diagram of the dynamics of the Lotka-Volterra equations, superposed with the contour plot of the log of the mean population in the limit $S \to \infty$, in the space $(\mu, \sigma)$ with $\gamma = 0$ and all carrying capacities equal to one. This plot is a reproduction using the equations derived in [Bunin, 2017, Roy et al., 2020]

.

mean abundance, this surface is used in Figure 4 of the main text for comparison between a purely random and the evolved interaction matrix.

### 2 First order perturbation theory of Lotka-Volterra equations

The equilibrium condition for the Lotka-Volterra equations is:

$$0 = N_i \left[ K_i - N_i - \sum_j \alpha_{ij} N_j \right] = N_i \left[ K_i - [(\mathbb{I} + \alpha)\mathbf{N}]_i \right] \tag{1}$$

We define $\chi_{ij} = \frac{\partial N_i}{\partial K_j}$ the perturbation matrix that measure the effect of a perturbation of the carrying capacities on the equilibrium abundances. We will denote with a $\star$ the vectors or matrices reduced to the set of extant species $\{i = 1, \ldots, S \,|\, N_i > 0\}$.

The solution of equation 1 is:

$$\begin{aligned}(\mathbb{I}^\star + \alpha^\star)\mathbf{N}^\star &= \mathbf{K}^\star \\ \mathbf{N}^\star &= (\mathbb{I}^\star + \alpha^\star)^{-1}\mathbf{K}^\star\end{aligned} \tag{2}$$

We now differentiate equation 1 with respect to $K_j$:

$$0 = \chi_{ij} \left[ K_i - N_i - \sum_k \alpha_{ik} N_k \right] + N_i \left[ \delta_{ij} - \chi_{ij} - \sum_k \alpha_{ik} \chi_{kj} \right] \tag{3}$$

For the set of extant species the first term is equal to zero and $N_i \neq 0$ so we must have :

$$\chi_{ij} + \sum_k \alpha_{i,k} \chi_{k,j} = \delta_{ij} \tag{4}$$

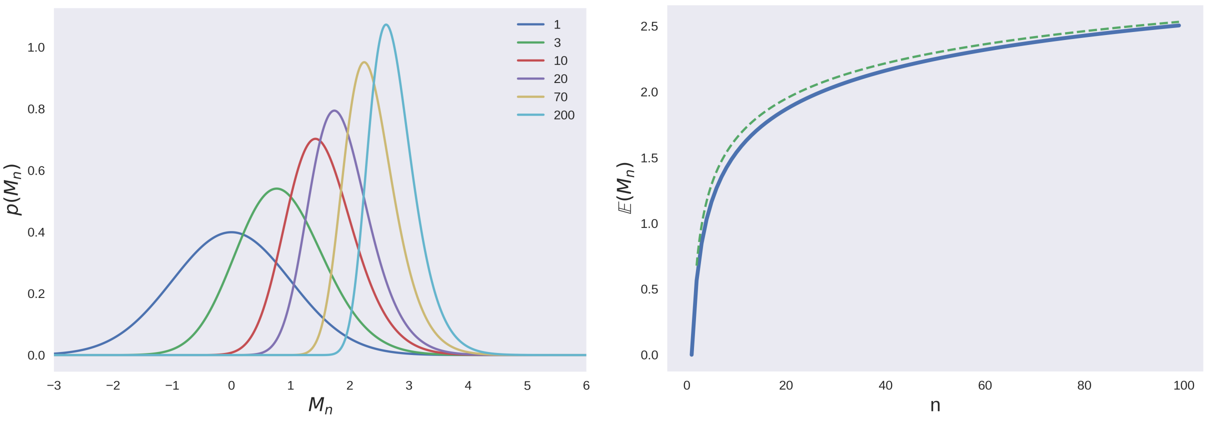


Figure 2: At the top is the distribution of $M_n$ for different values of n. At the bottom is the evolution of the expected value of this distribution with $n$ (in blue), with an approximation by $0.5 - \sqrt{2\log(n)}$ in dotted green.

That is to say, in matrix notation :

$$\chi^\star + \alpha^\star \chi^\star = \mathbb{I}^\star \tag{5}$$

This gives us an expression for $\chi^\star$:

$$\chi^\star = (\mathbb{I}^\star + \alpha^\star)^{-1} \tag{6}$$

and from equations 2 and 6 we get that:

$$\mathbf{N}^\star = \chi^\star \mathbf{K}^\star \tag{7}$$

### 3 Maximum over Gaussian samples

Let $\mathbf{x_1}, \ldots, \mathbf{x_n}$ be independent Gaussian random vectors of dimension $d$ and of law $\mathcal{N}(0, \mathbb{I}_d)$. For $\mathbf{u} \in \mathbb{R}^d$, we define $f_i = \mathbf{x_i} \cdot \mathbf{u}$. We want to find the distribution of $f_\star = \max(\{f_i\})$ and of the associated $\mathbf{x}_\star$.

We denote $\hat{\mathbf{u}} = \frac{\mathbf{u}}{\|\mathbf{u}\|}$. Let $P_u$ and $P_{u^\perp}$ be the projection matrices on $\hat{\mathbf{u}}$ and on its orthogonal space $\hat{\mathbf{u}}^\perp$ such that we have $f_i = \|\mathbf{u}\| P_u \mathbf{x_i}$. By Cochran theorem $P_u \mathbf{x_i}$ and $P_{u^\perp}\mathbf{x_i}$ are independent Gaussian variables of law $\mathcal{N}(0, P_u)$ and $\mathcal{N}(0, P_{u^\perp})$.

As $P_u \mathbf{x_i}$ is aligned with $\hat{u}$ and because $\hat{u}$ is normalized, we have $P_u \mathbf{x_i} = y_i \hat{\mathbf{u}}$ with $y_i \sim \mathcal{N}(0,1)$. We thus have $f_i = \|\mathbf{u}\| y_i$. In this notation $f_i > f_j \Leftrightarrow y_i > y_j$.

We define $M_n = \max(y_1, \ldots, y_n)$ such that $y_\star = M_n$ if $\forall i \neq \star,\, y_i < y_\star$ . By Bayes formula, $M_n$ has the probability density :

$$\begin{aligned} p(M_n = y) &= p(y_\star = y \,|\, \forall i \neq \star,\, y_i < y_\star) \\ &= \frac{1}{Z} \mathbb{P}(\forall i \neq \star,\, y_i < y) p(y_\star = y) \\ &= \frac{1}{Z} \left(\mathbb{P}(y_i < y)\right)^{n-1} p(y_\star = y) \\ &= \frac{1}{Z} \Phi(y)^{n-1} \phi(y) \end{aligned} \tag{8}$$

with $\Phi$ and $\phi$ the CDF and PDF of Gaussian law and $Z$ a normalization constant :

$$Z = \int_{-\infty}^{+\infty} \Phi(y)^{n-1}\phi(y)\,\mathrm{d}y = \frac{1}{n}\left[\Phi(y)^n\right]_{-\infty}^{+\infty} = \frac{1}{n} \tag{9}$$

Finally, $p(M_n = y) = n\Phi(y)^{n-1}\phi(y)$. This distribution is represented in Figure 2. It is asymptotic to a shifted Gumbel distribution [Gumbel, 2004].

We now use the decomposition $\mathbf{x}_\star = P_u \mathbf{x}_\star + P_{u^\perp}\mathbf{x}_\star = M_n \hat{\mathbf{u}} + P_{u^\perp}\mathbf{x}_\star$.

We thus have :

$$\mathbf{x}_\star = M_n \hat{\mathbf{u}} + \mathbf{b} \quad \text{with} \quad \mathbf{b} \sim \mathcal{N}(0, P_{u^\perp}) \tag{10}$$

or, in another notation :

$$\mathbf{x}_\star = M_n \hat{\mathbf{u}} + P_{u^\perp}\mathbf{z} = (M_n - \mathbf{z}\cdot\hat{\mathbf{u}})\hat{\mathbf{u}} + \mathbf{z} \quad \text{with} \quad \mathbf{z} \sim \mathcal{N}(0, \mathbb{I}_d) \tag{11}$$

### 4 Mutation-Selection Process for any $\gamma$

Let $\mathbf{N}$ be the equilibrium for the community described by the Lotka-Volterra equations. Each species' abundance obeys the equation:

$$0 = N_i \left[ K_i - N_i - \sum_j \alpha_{ij} N_j \right]. \tag{12}$$

After a mutational step, the interaction matrix becomes, at first order in $\varepsilon$ :

$$\hat{\alpha}_{ij} = \alpha_{ij} + \frac{\varepsilon\sigma}{\sqrt{S}} \eta_{ij} \tag{13}$$

with $\eta$ a Gaussian matrix of expected value zero, variance 1 and symmetric correlation $\gamma$.

Let $\hat{\mathbf{N}}$ be the equilibrium abundances associated to the interaction matrix $\hat{\alpha}$. For small $\varepsilon$, the difference $\delta\mathbf{N}$ between $\mathbf{N}$ and $\hat{\mathbf{N}}$ is of order $\varepsilon$. Then, to first order in $\varepsilon$, the equation for $\hat{\mathbf{N}}$ is:

$$0 = \hat{N}_i \left[ K_i - \hat{N}_i - \sum_j \alpha_{ij} \hat{N}_j \underbrace{- \varepsilon \frac{\sigma}{\sqrt{S}} \sum_j \eta_{ij} N_j}_{\delta K_i} \right] \tag{14}$$

The variable $\hat{\mathbf{N}}$ thus obeys, at equilibrium, equation 12, plus a perturbation field $\delta\mathbf{K} = -\varepsilon\frac{\sigma}{\sqrt{S}}\eta\mathbf{N}$ of order $\varepsilon$. Defining $\chi_{ij} = \frac{\partial N_i}{\partial K_j}$ as in section 2, for small $\varepsilon$ we can compute the first order perturbation $\delta\mathbf{N}$ as:

$$\begin{aligned} \delta\mathbf{N} &= \chi\delta\mathbf{K} \\ &= -\varepsilon\frac{\sigma}{\sqrt{S}}\chi\eta\mathbf{N} \end{aligned} \tag{15}$$

We now want to compute the induced variation $\delta f$ on the total abundance $f = \mathbf{1}\cdot\mathbf{N}$

$$\begin{aligned} \delta f &= \mathbf{1}\cdot\delta\mathbf{N} \\ &= -\varepsilon\frac{\sigma^t}{\sqrt{S}}\mathbf{1}^\top\chi\eta\mathbf{N} \\ &= -\varepsilon\frac{\sigma^t}{\sqrt{S}}\sum_{i,j,k} w_i\chi_{ij}\eta_{j,k}N_k \\ &= -\varepsilon\frac{\sigma^t}{\sqrt{S}}[(\chi^\top\mathbf{1})\otimes\mathbf{N}] : \eta \end{aligned} \tag{16}$$

With $\otimes$ the tensor product $((u\otimes v)_{ij} = u_i v_j)$ and $:$ the tensor contraction $(A : B = \sum_{ij} A_{ij}B_{ij})$ which is a scalar product in the space of matrices, thus allowing a generalization of the result from section 3 with matrices taken as vector of dimension $d = S^2$. However, in the case $\gamma \neq 0$, the $\eta_{ij}$ are not all independent. For that we use the decomposition of $\eta$ (see SI of [Barbier et al., 2018]) :

$$\eta = \frac{M + \kappa M^\top}{\sqrt{1+\kappa^2}} \tag{17}$$

with $\kappa = \frac{1-\sqrt{1-\gamma^2}}{\gamma}$ and $M$ a Gaussian matrix of mean 0, variance 1 with no correlations between $M_{ij}$ and $M_{ji}$.

Using $A : M^\top = A^\top : M$, we have

$$\delta f = -\varepsilon\frac{\sigma}{\sqrt{S}\sqrt{1+\kappa^2}}\left[(\chi^\top\mathbf{1})\otimes\mathbf{N} + \kappa\mathbf{N}\otimes(\chi^\top\mathbf{1})\right] : M \tag{18}$$

Now we can use the results from section 3. By substituting $\mathbf{x}$ with $M$ and $\mathbf{u}$ with $-\left[(\chi^\top\mathbf{1})\otimes\mathbf{N} + \kappa\mathbf{N}\otimes(\chi^\top\mathbf{1})\right]$, equation 11 yields:

$$\begin{aligned} M &= \frac{-M_n}{\mathcal{N}}\left[(\chi^\top\mathbf{1})\otimes\mathbf{N} + \kappa\mathbf{N}\otimes(\chi^\top\mathbf{1})\right] + B \\ \delta f &= \varepsilon\frac{\sigma\mathcal{N}}{\sqrt{S}\sqrt{1+\kappa^2}}M_n \end{aligned} \tag{19}$$

with $\mathcal{N} = \|(\chi^\top\mathbf{1})\otimes\mathbf{N} + \kappa\mathbf{N}\otimes(\chi^\top\mathbf{1})\|$ and $B$ a Gaussian matrix orthogonal (in terms of the scalar product $:$ ) to the selected "direction". We already get that the change in total abundance is :

$$N_T(\tau+1) - N_T(\tau) = \delta f = \frac{\varepsilon\sigma(\tau)M_n}{\sqrt{S}\sqrt{1+\kappa^2}}\sqrt{\sum_{ij}(v_iN_j + \kappa v_jN_i)^2} \tag{20}$$

that reduces to equation 6 of the main text when $\gamma = 0$. Putting the expression of $M$ in the formula for $\eta$ and using $\frac{2\kappa}{1+\kappa^2} = \gamma$ we get

$$\begin{aligned}\eta &= \frac{-M_n}{\mathcal{N}\sqrt{1+\kappa^2}} \left[(1+\kappa^2)(\chi^\top \mathbf{1}) \otimes \mathbf{N} + 2\kappa \mathbf{N} \otimes (\chi^\top \mathbf{1})\right] + \tilde{B} \\ &= \frac{-M_n\sqrt{1+\kappa^2}}{\mathcal{N}} \left[(\chi^\top \mathbf{1}) \otimes \mathbf{N} + \gamma \mathbf{N} \otimes (\chi^\top \mathbf{1})\right] + \tilde{B}\end{aligned} \tag{21}$$

with $\tilde{B} = \frac{B+\kappa B^\top}{\sqrt{1+\kappa^2}}$ being a Gaussian matrix of mean 0 and variance 1 with symmetric correlation $\gamma$ (it is the same construction as equation 17).

Putting this expression for $\eta$ in equation 13, we get :

$$\hat{\alpha}_{ij} = \alpha_{ij} - \frac{\varepsilon\sigma}{\sqrt{S}} \left( \frac{M_n\sqrt{1+\kappa^2}}{\mathcal{N}} \left[(\chi^\top \mathbf{1}) \otimes \mathbf{N} + \gamma \mathbf{N} \otimes (\chi^\top \mathbf{1})\right] - \tilde{B} \right) \tag{22}$$

For $\gamma = 0$, this yields equation (6) of the main text.

### 5 Correlation of the eigenvector with the abundances at equilibrium

Fig. 3 shows that at a late generation ($\tau = 2000$), the vector of abundances at equilibrium is strongly correlated with the eigenvector associated to the left-most isolated eigenvalue of $\alpha$ and to the vector $\mathbf{v} = \frac{\delta N_T}{\delta \mathbf{K}}$. Theses correlations explain the structure of $\alpha$: Fig. 4 shows that the species that are the most abundant are also the most mutualist.

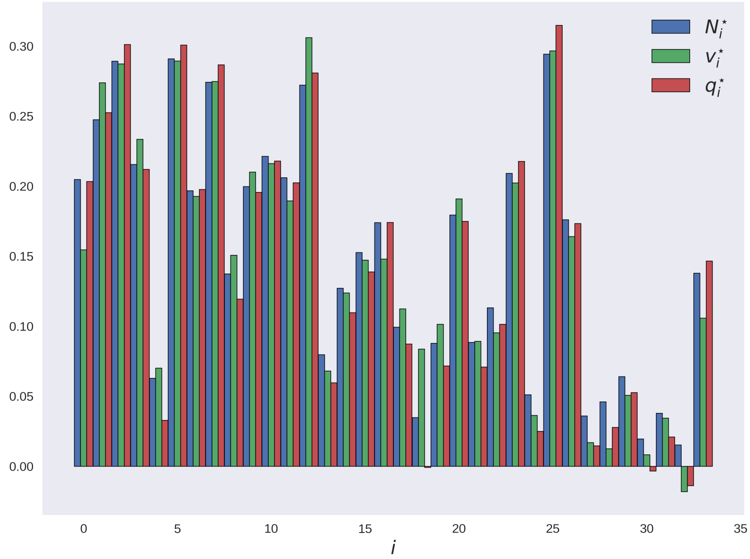


Figure 3: Coefficients of $\mathbf{N}^\star$, $\mathbf{v}^\star$ (as defined in the main text) and of the left eigenvector $\mathbf{q}^\star$ associated to the isolated eigenvalue of $\alpha^\star$ at generation $\tau = 2000$ reduced to extant species, all normalized, with indices sorted by decreasing carrying capacity. It is evident that $\mathbf{N}^\star$, $\mathbf{v}^\star$ and $\mathbf{q}^\star$ are strongly correlated. Their dependence on $\mathbf{K}$ is weaker, but still sufficient for the structure to emerge in Fig. 5 of the main text.

This correlation is caused by two effects that feed back onto one another. First, species with mutualistic interactions are more likely to be abundant than those with competitive interactions. Secondly, as shown in Eq. 7 (Main text), the most abundant species will see their interactions evolve faster toward mutualism.

### 6 Limit of small $\sigma$ and large $S$

For simplicity, we consider here that all the carrying capacities are equal (we set $K_i = 1$, without loss of generality). When $S \gg 1$, $\sigma(\tau)$ is constant, and we assume here that it is initially small.

For small $\sigma$, the interaction matrix $\alpha(\tau)$ can be characterized only by its mean value :

$$\alpha_{ij}(\tau) = \frac{\mu(\tau)}{S} + \mathcal{O}(\sigma) \tag{23}$$

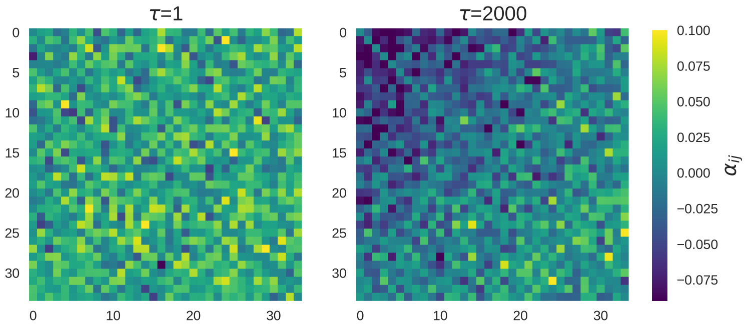


Figure 4: Coefficients of the interaction matrix $\alpha$ at generations 1 (left) and 2000 (right) with rows and columns sorted by decreasing **equilibrium abundances at** $\tau = 2000$ for the same simulation as described in Materials and Methods. Only the species that have positive abundance at generation 2000 are shown.

that we write in matrix notation :

$$\alpha(\tau) = \frac{\mu(\tau)}{S}\mathbf{1}\mathbf{1}^\top + \mathcal{O}(\sigma) \tag{24}$$

Then, following eq 6, the perturbation matrix can be expressed as :

$$\begin{aligned} \chi(\tau) &= (\mathbb{I} + \alpha(\tau))^{-1} \\ &= \left(\mathbb{I} + \frac{\mu(\tau)}{S}\mathbf{1}\mathbf{1}^\top + \mathcal{O}(\sigma)\right)^{-1} \\ &= \mathbb{I} - \frac{\mu(\tau)/S}{1+\mu(\tau)}\mathbf{1}\mathbf{1}^\top + \mathcal{O}(\sigma) \end{aligned} \tag{25}$$

using Sherman–Morrison formula. With this, we can compute the abundances at equilibrium:

$$\begin{aligned} \mathbf{N}(\tau) &= \chi(\tau)\mathbf{1} \\ &= \frac{1}{1+\mu(\tau)}\mathbf{1} + \mathcal{O}(\sigma) \end{aligned} \tag{26}$$

The total abundance along the evolutionary trajectory depends, as well as the interaction matrix, only on the mean interaction strength:

$$\frac{N_T(\tau)}{S} = \langle N_i(\tau)\rangle = \frac{1}{1+\mu(\tau)} + \mathcal{O}(\sigma). \tag{27}$$

In the same fashion, we get that :

$$\mathbf{v}(\tau) = \frac{1}{1+\mu(\tau)}\mathbf{1} + \mathcal{O}(\sigma) \tag{28}$$

Equation 22 for $\gamma = 0$ or equation (6) of the main text then gives the recursive equation for $\mu(\tau)$:

$$\mu(\tau+1) = \mu(\tau) - \sqrt{1+\gamma}\frac{\varepsilon M_n \sigma}{\sqrt{S}} \tag{29}$$

that has for solution:

$$\mu(\tau) = \mu(0) - \tau \cdot \frac{\varepsilon\sigma}{\sqrt{S}}\overline{M_n}\sqrt{1+\gamma} \tag{30}$$

Fig. 5 shows that equations 27 and 30 match the numerical simulations remarkably well in the case of an initial $\sigma = 0.1$ and $S = 100$.

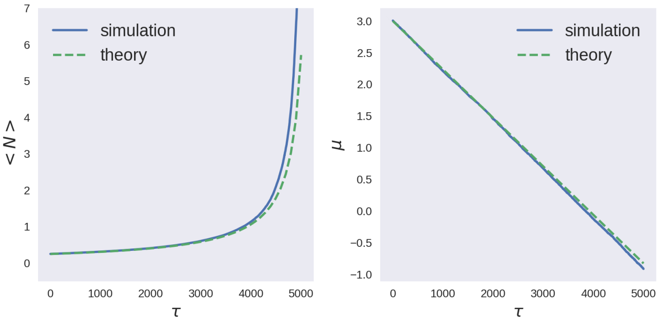


Figure 5: Comparison of the evolution of the mean abundance $< N >$ and of the rescaled mean interaction $\mu$ from the Eqs. 30 and 27 (green lines) and from numerical simulations (blue lines) with $S = 100$ and $\sigma(0) = 0.1$.

### 7 Community changes in the absence of selection

We now look at the evolutionary dynamics of the interaction matrix in the neutral regime, when any community leaves an offspring with equal probability.

In this subsection only, we denote for simplicity $d = S(S-1)$ the number of interaction terms and every sum $\sum$ are index on $i \neq j$. The notations $b_{ij}$ is not the same as before.

The process is the following:

- We have $\alpha_{ij}^{(t)} = m_t + \mathrm{std}_t a_{ij}^{(t)}$ with $m_t = \frac{1}{d}\sum \alpha_{ij}^{(t)}$ the empirical mean and $\mathrm{std}_t = \sqrt{\frac{1}{d}\sum(\alpha_{ij}^{(t)} - m_t)^2}$ the empirical standard deviation.
- We then define $\alpha_{ij}^{(t+1)} = m_t + \mathrm{std}_t b_{ij}^{(t+1)}$ with $b_{ij}^{(t+1)} = \frac{a_{ij}^{(t)} + \varepsilon \eta_{ij}^{(t+1)}}{\sqrt{1+\varepsilon^2}}$.
- We repeat the operation.

Note that by definition, $a_{ij}$ have the following properties: $\sum a_{ij}^{(t)} = 0$ and $\frac{1}{d}\sum(a_{ij}^{(t)})^2 = 1$.

We now wish to get a an expression of $\mu_{t+1}$ as a function of $\mu_t$ and $\eta$. We start with the empirical mean:

$$\begin{aligned} m_{t+1} &= \frac{1}{d}\sum \alpha_{ij}^{(t+1)} \\ &= m_t + \frac{\mathrm{std}_t}{d}\sum b_{ij}^{(t+1)} \\ &= m_t + \frac{\mathrm{std}_t\,\varepsilon}{\sqrt{1+\varepsilon^2}}\frac{1}{d}\sum \eta_{ij}^{(t+1)} \end{aligned} \tag{31}$$

Using the Central Limit Theorem and writing with $\mu$, $S$ and $\sigma$ we get:

$$\boxed{\mu_{t+1} \sim \mathcal{N}(\mu_t, \sigma_t \frac{\varepsilon}{\sqrt{S(1+\varepsilon^2)}})} \tag{32}$$

To have a similar expression for $\sigma$ the computation is a bit longer. We start with the empirical variance:

$$\begin{aligned} \mathrm{Var}_{t+1} &= \frac{1}{d}\sum(\alpha_{ij}^{(t+1)} - m_{t+1})^2 \\ &= \frac{1}{d}\sum(m_t - m_{t+1} + \mathrm{std}_t b_{ij}^{(t+1)})^2 \\ &= \frac{\mathrm{Var}_t}{d}\sum\left(\frac{a_{ij}^{(t)} + \varepsilon\eta_{ij}^{(t+1)}}{\sqrt{1+\varepsilon^2}} - \frac{\mathrm{std}_t\,\varepsilon}{\sqrt{1+\varepsilon^2}}\frac{1}{N}\sum_{k\neq l}\eta_{k,l}^{(t+1)}\right) \end{aligned}$$

We denote $\langle\eta\rangle = \frac{1}{d}\sum_{k\neq l}\eta_{k,l}^{(t+1)}$ the empirical mean of $\eta$ so that:

$$\begin{aligned} \mathrm{Var}_{t+1} &= \frac{\mathrm{Var}_t}{(1+\varepsilon)^2 d}\sum\left(\alpha_{ij}^t + \varepsilon(\eta_{ij} - \langle\eta\rangle)\right) \\ &= \frac{\mathrm{Var}_t}{(1+\varepsilon)^2 d}\left[\sum(a_{ij}^t)^2 + \varepsilon\sum(\eta_{ij} - \langle\eta\rangle)^2 + 2\varepsilon\sum a_{ij}^{(t)}(\eta_{ij} - \langle\eta\rangle)\right] \end{aligned}$$

We now use the fact that $\sum a_{ij}^{(t)} = 0$ and $\frac{1}{d}\sum(a_{ij}^{(t)})^2 = 1$. Furthermore, as $\eta$ is a Gaussian random variable, $\sum(\eta_{ij} - \langle\eta\rangle)^2 \sim \chi_{n-1}^2$ (chi-square distribution).

As $a_{ij}$ and $\eta_{ij}$ are uncorrelated we have by the Central Limit Theorem:

$$\frac{1}{d}\sum a_{ij}\eta_{ij} \sim \mathcal{N}(0, \frac{1}{\sqrt{N}}) \tag{33}$$

We then use the asymptotic expression: $\chi^2_{d-1} \longrightarrow d-1+\sqrt{2(d-1)}\mathcal{N}(0,1)$ so that in the large $S$ (and thus $d$) limit:

$$\mathrm{Var}_{t+1} = \frac{\mathrm{Var}_t}{(1+\varepsilon)^2}\left[1+\varepsilon^2+\varepsilon^2\sqrt{\frac{2}{d}}\mathcal{N}(0,1)+\frac{2\varepsilon}{\sqrt{d}}\mathcal{N}(0,1)\right]$$

But as $\varepsilon \ll 1$ we can neglect the first normal distribution:

$$\mathrm{Var}_{t+1} = \mathrm{Var}_t\left(1+\frac{2\varepsilon}{1+\varepsilon^2}\frac{1}{\sqrt{d}}\mathcal{N}(0,1)\right) \tag{34}$$

Using the Taylor expansion of the square root we get the expression for $\sigma$:

$$\boxed{\sigma_{t+1} \sim \mathcal{N}(\sigma_t, \sigma_t\frac{\varepsilon}{S(1+\varepsilon^2)})} \tag{35}$$

We thus observe that this process is a isotropic random walk in the space $(\mu, \sigma)$ and that the variation in $\sigma$ are small compared to the one in $\mu$: it scales like $\frac{1}{S}$ for $\sigma$ and $\frac{1}{\sqrt{S}}$ for $\mu$. Thus, for large $S$, $\sigma$ evolves with a longer time-scale than $\mu$ and so can be approximated as constant. These results are consistent with the numerical observation in Figure 6.

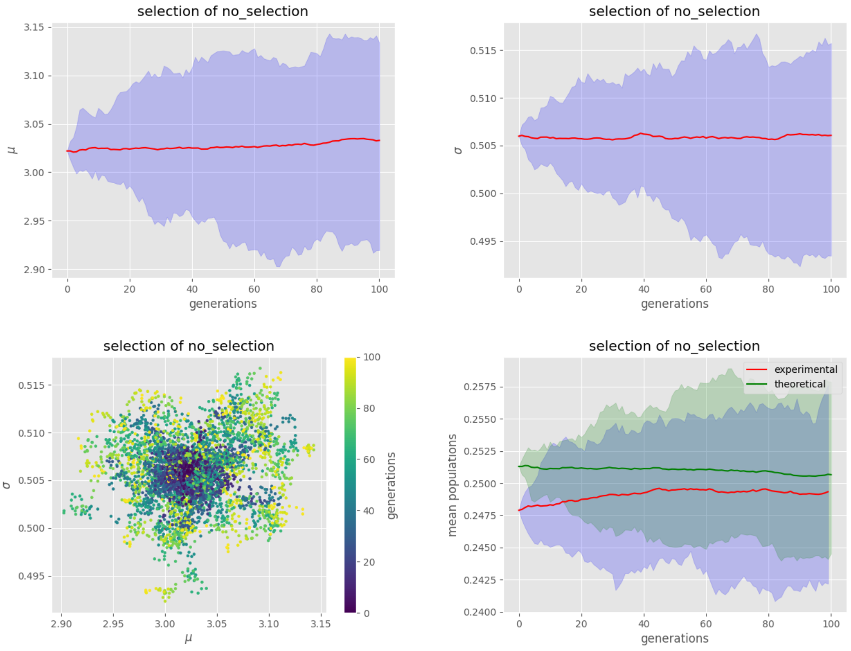


Figure 6: Evolution of the empirical $\mu$ (top-left), $\sigma$ (top-right) and mean population $\langle N\rangle$ (bottom-right) in absence of selection, with the parameters $S = 100$, $\varepsilon = 0.1$, $\tau_{\max} = 100$, $n = 50$ and $\gamma = 0$. For the panels in the top and in the bottom-right, the red lines are the mean of the quantity of interest and the blue area is bounded by the maximum and minimum values. The green line in the bottom-right panel represent the predicted value from the cavity method. The panel in the bottom-left is the evolution of the communities in the $(\mu, \sigma)$ space.

### 8 Evolution of species diversity

Fig. 7 represents the evolution of the diversity (or species richness) $\phi = \frac{S^\star}{S}$ along the same evolutionary trajectory as the figures of the main text. This shows that diversity decreases along the evolutionary trajectory but the number of extant species does not collapse, and the community maintains a substantial amount of diversity.

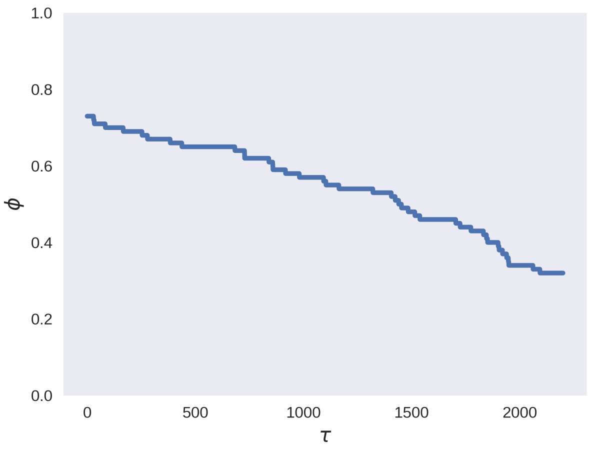


Figure 7: **Changes of species richness along an evolutionary trajectory.** Selection for increased total abundance leads to a decrease in the species richness. The parameters of the simulation are described in the Methods section.

### 9 Comparison with synthetic interaction matrix

Following [Barbier et al., 2021], we can compute the inferred interaction matrix $\beta$ that gives rise to the observed equilibrium abundances and has a given mean interaction $\bar{\beta}$. Neglecting the correlations between different matrix elements (which are sub-leading in the large $S$ limit), such matrix is of the form:

$$\beta^\star_{ij} = \bar{\beta} + \frac{(K_i - N_i - \bar{\beta})N_j}{\sum_k N_k^2} + \sigma B_{ij} \tag{36}$$

with $B$ a Gaussian matrix of mean 0 and variance 1. Fig. 8 shows the eigenvalue distribution of such matrix, compared to the one obtained from the evolutionary process. The agreement is remarkable.

### 10 Generalizations

We checked that the features presented in the main text hold for a broad range of parameters in the numerical simulations, in particular for most initial values of $(\mu, \sigma)$ as long as we are in the unique equilibrium phase (see section 1), for any values of $m$ and $n$ (when selecting $m$ communities out of $n$) as long as $n > 1$ and $m < n$ (theses two extreme cases lead to no selection) and for any value of $\gamma$ with the exception of $\gamma = -1$, as expected from the equations discussed in the main text. The addition of a small immigration term to the ecological dynamics, moreover, doesn't qualitatively alter the results.

We considered different selection functions, such as $f = \sum_i w_i N_i$ with all weights $w_i$ being positives. Similar results keep holding. This can be understood by interpreting the selection process as a modification of selection for the total abundance, obtained by rescaling species abundances as $w_i N_i$, so that species $i$ has carrying capacities $w_i K_i$ and an interaction matrix $\alpha_{ij} w_i / w_j$. The same computations explained in the main text and in Section 4 then allow us to obtain the exact same recursive equations for $f$ and $\alpha$ but with $\mathbf{v} = \frac{\delta f}{\delta K}$ and $\mathbf{v}^\star(\tau) = (\mathbb{I}^\star + \alpha^\star(\tau)^\top)^{-1}\mathbf{w}^\star$.

### 11 Numerical integration method

We here present an integration method for the Lotka-Volterra equations :

$$\frac{\mathrm{d}N_i}{\mathrm{d}t} = r_i \frac{N_i}{K_i}\left(K_i - N_i - \sum_{j \neq i} \alpha_{ij} N_j\right), \tag{37}$$

with $K_i$ the carrying capacities, $r_i$ the growth rates (all equals to one in the paper) and $\alpha$ the interaction matrix.

In contrast to the Euler method where one assumes that the derivative is constant during a short interval of time $\mathrm{d}t$, potentially leading to negative populations for strong derivatives, we assume that only the abundances of the other species $(N_j(t))_{j \neq i}$ are constant. During this time interval $[t, t + \mathrm{d}t]$, the equations are reduced to uncoupled logistic equations of effective carrying capacities $\widetilde{K_i} = K_i - \sum_{j \neq i} \alpha_{ij} N_j(t)$ and effective growth

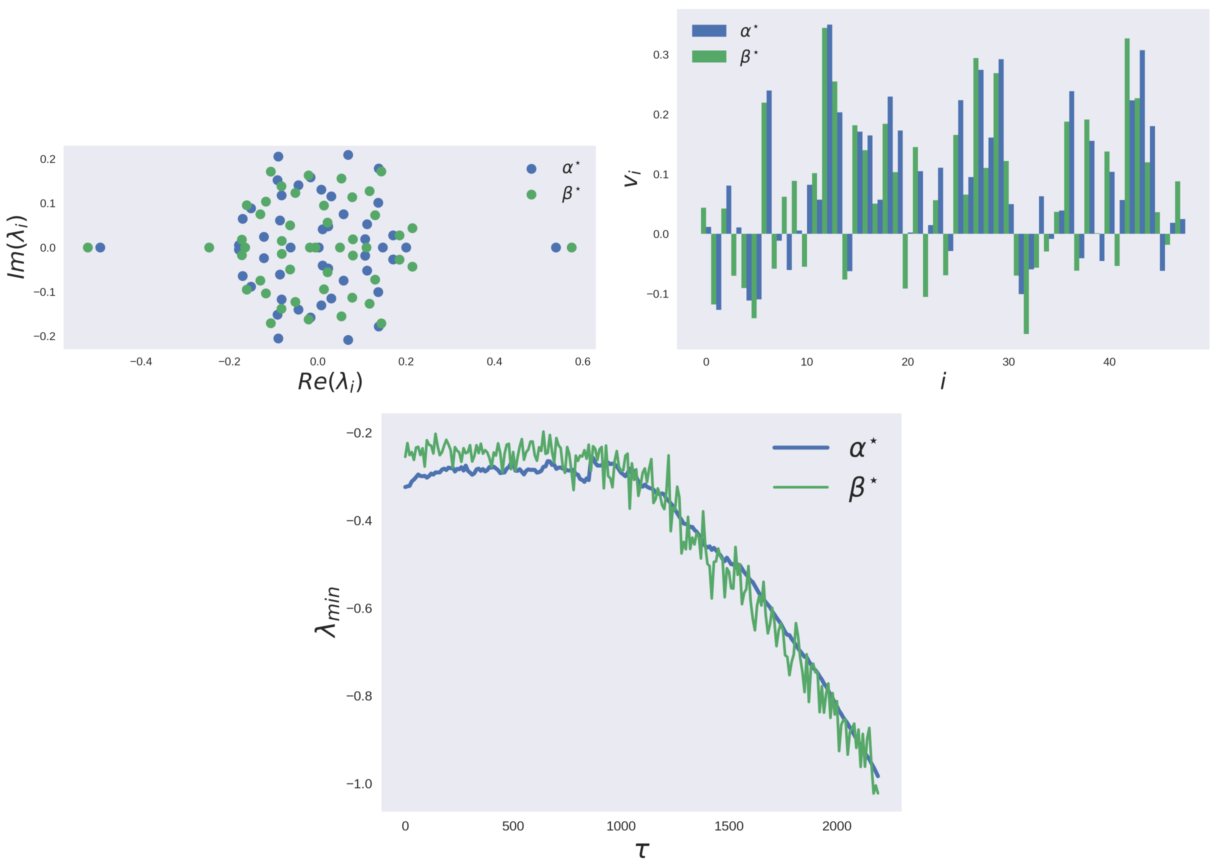


Figure 8: **Evolved and synthetic matrix have similar structure of eigenstates.** Comparison between the eigenvalues of the evolved interaction matrix $\alpha^\star$ (blue) and of the maximum entropy matrix $\beta^\star$ (green) at generation $\tau = 1500$ (Top) and the coefficient of the eigenvector of the minimal eigenvalue (middle). Evolution of the minimum eigenvalue of both matrices for every generations (Bottom).

rates $\widetilde{r}_i(t) = r_i \widetilde{K}_i(t)/K_i$. These logistic equations can be analytically solved in the interval $[t, t + \mathrm{d}t]$, giving the recursive formula:

$$N_i(t + \mathrm{d}t) = \frac{N_i(t)\widetilde{K}_i(t)}{N_i(t) + (\widetilde{K}_i(t) - N_i(t))\exp\Big(-\widetilde{r}_i(t)\mathrm{d}t\Big)} \tag{38}$$

with

$$\begin{cases} \widetilde{K}_i(t) = K_i - \sum_{j\neq i} \alpha_{ij} N_j(t) \\ \widetilde{r}_i(t) = r_i \frac{\widetilde{K}_i(t)}{K_i} \end{cases} \tag{39}$$

We then have a logistic-by-part curve that avoids abundances to become negative. We can show by Taylor expansion that we get back the Euler method at first order in $\mathrm{d}t$.

As we can see in Figure 9, this logistic integration scheme is more stable when we increase the time step.

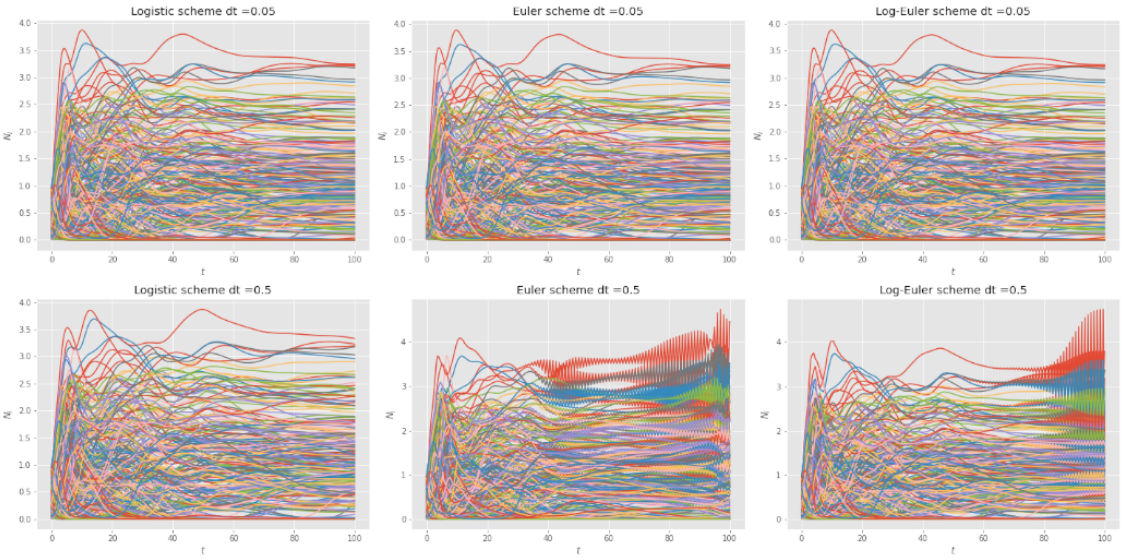


Figure 9: **Comparison of different integration schemes with different time steps** $dt$**.** On the left is the Logistic scheme presented here, in the middle the classic Euler method and on the right the Euler method for the logarithm of $N_i$. The three on top are for a time-step $dt = 0.05$ and the three below are for $dt = 0.5$. We consider here a trajectory in the chaotic regime (parameters $S = 500$, $\mu = 2$, $\sigma = 1.5$), where simulations are most problematic, as the trajectory approaches the boundary of the phase space.
